# Effects of Escitalopram and Relearning on Cortical and Subcortical Grey Matter in Healthy Humans

**DOI:** 10.1101/2021.04.24.440965

**Authors:** T Vanicek, MB Reed, R Seiger, M Godbersen, M Klöbl, J Unterholzner, B Spurny, G Gryglewski, P Handschuh, C Kraus, T Stimpfl, R Rupprecht, S Kasper, R Lanzenberger

## Abstract

The antidepressant effect of selective serotonin reuptake inhibitors (SSRI) is related to increased neuroplasticity during relearning. Stress-induced dendritic atrophy in key brain areas for learning and memory such as the hippocampus and prefrontal cortex is reversed by SSRI treatment. This finding is accompanied by behavioral stabilization. The aim of this study was to investigated serotonergic modulation effects on structural neuroplasticity (cortical thickness, subcortical volumes) during relearning in healthy subjects. Participants performed daily associative learning tasks over 3 weeks followed by a 3-week relearning phase combined with intake of the SSRI escitalopram or placebo. Evidence suggests that SSRIs promote the brains susceptibility to change on the basis of environment factors. We found no effect of SSRI on grey matter measures during relearning. Here, non-findings might be a consequence of the implemented intensity and duration of study interventions. With sparse literature on healthy participants in this field, future studies will have to further elucidate SSRIs properties on relearning and structural neuroplasticity.

## Introduction

The brain’s capacity to adapt to a changing environment and meet its needs in order to navigate effectively past life challenges is grounded in its plasticity. Synaptic (re)organization, dendritic remodeling, neuron-glial coupling and epigenetic processes are mechanisms that underlie neuroplasticity and consequently represent the fundament of memory formation and consolidation (Bruel-Jungerman, et al., 2007; Draganski, et al., 2004; Forrest, et al., 2018; Matsuzaki, et al., 2004; Sudhof, 2018). Brain activity-dependent pre- and postsynaptic specialization peaks in adolescence and continues, at significantly lower turnover rates, throughout life-time (Forrest, et al., 2018; Sudhof, 2018).

Serotonergic modulation has been found to influence learning and memory performances across species, as the depletion of tryptophan (a precursor of serotonin) results in diminished cognitive performance in animals, healthy subjects and patients (Clarke, et al., 2005; Neumeister, et al., 2004; Riedel, et al., 1999). In a systematic review with a focus on the effects of chronic SSRI administration in healthy subjects, Knorr et al. reported significant changes to various physiological, behavioral and neurophysiological parameters (Knorr, et al., 2019). Moreover, memory consolidation and relearning were improved by SSRIs in Long-Evans rats (Brown, et al., 2012).

Among various molecular pathways involved in neuronal organization and adaptation processes, the serotonergic neurotransmitter system affects neuronal circuit formation by regulating neuroplastic processes at the synapse level (Dayer, 2014). Moreover, serotonergic agents such as selective serotonin reuptake inhibitors (SSRIs) are frequently prescribed for irregular mental states such as depression and anxiety (Bauer, et al., 2013). These agents have been found to unfold efficacy by modulating cell cascades relevant for neuronal restructuring. SSRIs elevate protein synthesis of the cyclic adenosine monophosphate (cAMP) response element binding protein (CREB) and brain derived neurotrophic factors (BDNF). These proteins are relevant for synaptic formation and hence, for memory and learning (Nibuya, et al., 1995; Park and Poo, 2013; Simmons, et al., 2009). Recent discoveries by Castrén and his group point towards a direct link between SSRIs and neuroplasticity that is partly ascribed to their binding properties to neurotropic growth factor receptors (Casarotto, et al., 2021). Animal studies using stress models to explore the relationship of neuropathology and behavior demonstrate an association between dendritic degeneration and cognitive domains, such as impaired working memory and attentional set-shifting deficiencies (Cerqueira, et al., 2007; Liston, et al., 2006). Noticeably, SSRIs have been shown to reverse both stress-induced hippocampal and prefrontal dendritic atrophy as well as depressive behavior in rodents. Furthermore, it has been discovered that SSRIs contribute to balanced neurotransmission through enhancing the synthesis of synaptic cell adhesion molecules (Bessa, et al., 2009; Hajszan, et al., 2009).

Within the field of neuroimaging, several studies focused on the effects of SSRIs on neural activation during emotional and neutral processing using functional magnetic resonance imaging (MRI) (Di Simplicio, et al., 2014; Norbury, et al., 2009; Sladky, et al., 2015). However, very few studies address structural brain changes after long-term SSRI administration. In two imaging studies, SSRIs were found to affect grey matter in healthy humans and in non-human primates, with findings suggesting divergent SSRI effects in healthy compared to depressive subjects (Kraus, et al., 2014; Shively, et al., 2017). Further investigations are needed, since the mechanisms underlying serotonergic modulation of grey matter adaptations in combination with learning and memory consolidation are poorly understood.

We conducted a longitudinal SSRI- and learning-intervention study to test the effects of serotonergic modulation on neuroplasticity using structural MRI during reversal learning. The aim of this study was to explore the neurobiological effects of SSRI on relearning as a model for treatment properties of SSRIs in neuropsychiatric disorders with relations to cognitive flexibility and neuronal reorganization. We hypothesized a SSRI effect on brain areas comprising the hippocampus and parahippocampus during reversal learning, whereas emotional content learning would expand recruitment of emotion regulating regions as the amygdala, prefrontal cortex, insular and the anterior cingulate cortex. Healthy subjects performed an associative learning task with and without emotional content for 3 weeks, had to perform relearning within groups for additional 3 weeks, whereas received escitalopram 10 mg or placebo during the second period.

## Methods

### Study design

To assess SSRI-invoked structural brain changes, we conducted a randomized placebo-controlled and double-blind (re)learning study with healthy participants. Structural MRI measurements were carried out at 3 time points throughout the study: at baseline, after associative learning and finally after associative relearning, with a respective time-interval of 3 weeks between sessions. Before MRI-assessment and associative learning, subjects were randomly assigned to an associative learning group with or without emotional valence as well as to the substance group (escitalopram vs. placebo). The emotional group consisted of pairwise matching of face pictures, whereas the group without emotional content comprised of associations of Chinese characters with unrelated German nouns. After a 3-week learning period, subjects either receiving a daily dose of 10 mg escitalopram or placebo while relearning new associations of the previously learned ones for a further 3 weeks. For details and illustration see Figure 1.

**Figure 1.**
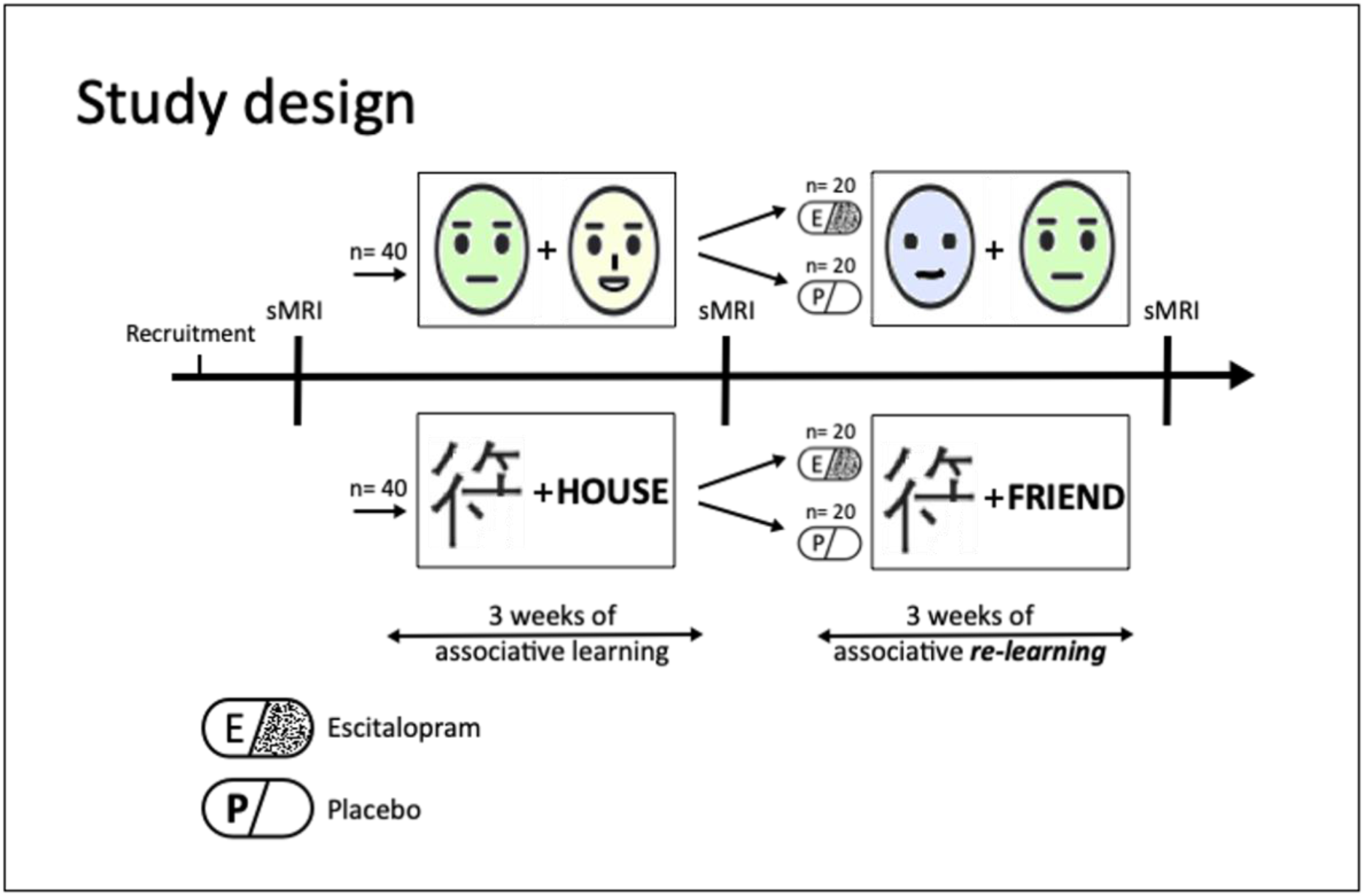
Study design. Healthy subjects performed an association learning paradigm with (face pairs; note: to meet bioRxiv policy, photographs of faces in this figure are replaced by images) and without emotional content (Chinese characters and unrelated German nouns) on a daily base for three weeks. Subsequently, subjects had to relearn shuffled associations (within their group) for a further 3 weeks while receiving either a daily dose of 10 mg escitalopram or placebo. MRI measurements were scheduled at baseline, after associational learning (3 weeks) and after associational relearning (another 3 weeks). Study subjects were randomly assigned to learning groups and treatment conditions (escitalopram vs. placebo). Abbreviations: sMRI structural magnetic resonance imaging.

### Subjects

The planned sample size of this study was 80 healthy subjects that completed the study. General health was assessed through the medical history and a physical examination. A structured clinical interview for DSM-IV (SCID I) was conducted to assure mental health. Besides general health, willingness and competence to partake in this study, further inclusion criteria consisted of being between 18 and 55 years of age, right-handed and non-smoking. Exclusion criteria comprised any medical, psychiatric or neurological illness, any lifetime use of SSRIs, first-degree relatives with a history of psychiatric illness, color blindness, non-European ancestry, MRI contraindications and knowledge of the Japanese Kanji or the Chinese Hanzi. Subjects that dropped out due to any reason(s) listed above were replaced.

All participants gave written consent to partake in this study and received reimbursement for their participation. The study was approved by the ethics committee of the Medical University of Vienna (EK Nr.: 1739/2016) and was performed in accordance with the Declaration of Helsinki (1964). The study was registered at clinicaltrials.gov with the identifier NCT02753738. The distributions of sex and age in between groups were tested using Fisher’s exact and that of age with a Kruskal-Wallis test.

### Associative learning paradigm

Each subject was required to learn 200 image pairs via an online platform, which was developed in-house. The image pairs were presented sequentially for 5 seconds, whereas each session consisted of a pseudorandom subselection (sampling with replacement) of 52 image pairs. Each learning session was followed by a retrieval phase with a pseudorandom selection of 52 single images from all previously learned pairs. The correct association had to be selected out of 4 possible answers without time limit. The faces were extracted from the “10k Adult Faces Database” (Bainbridge, et al., 2013). The Chinese characters were randomly selected and had no connection to the associated German nouns. Learning days 1, 22 and 42 were completed during MRI session.

### Study Drug Administration and Monitoring

The verum subgroups received Escitalopram (Cipralex® Lundbeck A/S, provided by the Pharmaceutical Department of the Medical University of Vienna) 10 mg orally per day for 21 days during relearning (i.e., after the 2^nd^ MRI). The other half of the study subjects received placebo tablets for 21 days as a control group. Venous blood was drawn from the cubital vein to assess citalopram plasma through levels at 3 time points: 1, 2 and 3 weeks after therapy start (the last blood sampling was performed directly before the 3^rd^ MRI). Citalopram plasma levels were assessed with liquid chromatography–tandem mass spectrometry (LC-MS/MS) at the Clinical Department of Laboratory Medicine of the Medical University of Vienna. According to the literature the therapeutic reference range for escitalopram it is 15-80 ng/mL (Hiemke, et al., 2011).

### MRI acquisition and Processing

Each MRI session was conducted using a 3 Tesla MR Scanner (MAGNETOM Prisma, Siemens Medical, Erlangen, Germany) and a 64-channel head coil at the High-field MR Center, Medical University of Vienna. Whole brain T1 images were acquired during each MRI session, (Repetition time (TR) = 2300 ms; echo time (TE) = 2.95 ms; inversion time (TI) = 900 ms; flip angle (α) = 9°; PAT = GRAPPA2; matrix = 240 x 256, 176 slices; 1.05 x 1.05 x 1.20 mm3; acquisition time (TA) = 5:09 min)

### Surface-based analysis using FreeSurfer 6.0

The automated recon-all pipeline implemented in the FreeSurfer 6·0 software with default parameters was used for cortical surface reconstruction and parcellation of 34 cortical regions for each hemisphere and 9 subcortical regions (Harvard Medical School, Boston, USA; http://surfer.nmr.mgh.harvard.edu/). A within-participant template was created via inverse consistent registration using all time points for the longitudinal processing pipeline, which included the following processing steps: skull stripping, Talairach registration, and initialization of cortical surface reconstruction, cortical atlas registration, and subcortical parcellation (Saygin, et al., 2017). Cortical thickness as well as subcortical volumes for the Desikan-Killiany (DK) atlas (Desikan, et al., 2006) were extracted at each time point. After the automated processing, all volumes were visually inspected.

### Voxel-based morphometry (VBM) analysis using CAT12

To calculate the voxel-based morphometric changes, data was processed using the CAT12 toolbox (http://www.fil.ion.ucl.ac.uk/spm/software/spm12/,version7771) in MATLAB (version 9.4) via the CAT12 longitudinal pipeline with default settings. Prior to preprocessing, all raw data were visually inspected for potential artifacts. For each participant, the scans for all 3 time points were registered, resampled and bias-corrected. Each scan was then skull stripped and segmented into grey matter, white matter and cerebrospinal fluid. Finally, these maps were transformed into MNI space and spatially smoothed using an 8-mm Gaussian kernel. Lastly, the CAT12 reports and output data were visually controlled for miss registrations and segmentation errors.

### Statistics

The effects of substance, learning content and time on cortical thickness and subcortical volumes were analyzed using SPSS 25.0. To this end, four-way repeated measures analyses of variance (rmANOVAa) were set up to test for substance x content x time x region of interest (ROI) interaction effects. rmANOVAs were performed separately for cortical and subcortical regions.

Statistical analysis of the VBM data was performed using the CAT12 toolbox within SPM12 (https://www.fil.ion.ucl.ac.uk/spm/). To elucidate substance x content x time interaction effects on grey matter a 2 x 2 x 3 repeated measures ANOVA was modelled. VBM analyses were corrected for multiple testing using Gaussian random field theory as implemented in SPM12 and the threshold for significance was set at P⩽0.05 family-wise error (FWE)-corrected at the cluster-level following P⩽0.001 uncorrected at the voxel-level. Interaction effects were dropped in case of non-significance in both surface-based and VBM analyses. Bonferroni correction was used to correct for all post-hoc comparisons in the VBM analysis and for the number of ROIs and post-hoc tests for the FreeSurfer parcellation. To correct for different brain sizes and volumes the total intracranial volume was included as a covariate.

## Results

### Study population

Out of the 138 subjects recruited only 84 successfully completed both phases of the study. Of the 84 subjects, 4 were dropped due to insufficient data quality, further 3 subjects had to be excluded as their escitalopram plasma levels were under the measurable threshold (< 10 ng/mL) and 1 subject was dropped due to irregular learning performance. For the analyses, 76 subjects were encompassed the final sample which included, 44 women, 32 men, with a mean age (±SD) of 25.6 ± 5 years. No significant group differences regarding age, sex or group participation numbers (p > 0.15) were found. See table 1 for detailed demographics and participant stratification.

**Table 1:** Demographics for all participants included in the final analysis. (A) indicates the demographics of the 1st 3-week learning phase, were subjects were assigned to either the faces or character subgroup. (B) shows the demographic data for the 2nd phase, where subjects had to relearn previously learnt associations while taking 10 mg/day escitalopram or placebo.

| (A) | Faces |  |  |  | Character |  |  |  |
| --- | --- | --- | --- | --- | --- | --- | --- | --- |
|  | Mean |  | SD |  | Mean |  | SD |  |
| N | 38 |  |  |  | 38 |  |  |  |
| Age [years] | 25.61 |  | 5.40 |  | 25.39 |  | 4.48 |  |
| Gender [M/F] | 13/25 |  |  |  | 19/19 |  |  |  |
| (B) | Verum |  | Placebo |  | Verum |  | Placebo |  |
|  | Mean | SD | Mean | SD | Mean | SD | Mean | SD |
| N | 16 |  | 22 |  | 19 |  | 19 |  |
| Age [years] | 25.50 | 5.21 | 25.91 |  | 25.53 | 4.94 | 25.26 | 4.10 |
| Citalopram plasma concentration [ng/mL] | 18.01 | 10.51 | 0 | 0 | 18.97 | 9.28 | 0 | 0 |
| Gender [M/F] | 7/9 |  | 6/16 |  | 8/11 |  | 11/8 |  |

### Analysis of grey matter volume and thickness

To test our hypothesis of SSRIs effecting the (para)hippocampal structures during relearning and emotional content learning increasing the involvement of the amygdala, prefrontal cortex, insular and the anterior cingulate cortex, a two-way rmANOVA was conducted to analyze interaction effects on the brain regions listed above. To this end, 2 separate two-way rmANOVA models were conducted to analyze substance, content by time and ROI interactions. We found no significant 4-way or lower interactions in both cortical and subcortical regions (P_Bonferroni_ > 0.05).

To test for interaction effects outside of our hypothesis on the remaining cortical and subcortical parcels, further 2 two-way rmANOVAs were run. No significant interactions (substance, content by time, ROI) in either cortical or subcortical structures were discovered (P_Bonferroni_ > 0.05).

### Analysis of voxel-based grey matter changes

To assess the influence of substance and learning content interactions on cortical gray matter and subcortical volume, a VBM analysis was also conducted. Again, no 3-way or lower interactions (substance, content by time) were found after correction for multiple comparisons P_Bonferroni_ > 0.05.

## Discussion

In this study, we aimed to evaluate the long-term effects SSRI administration in combination with relearning had on grey matter in healthy humans *in vivo*. Thus, subjects were enrolled in an associative learning study having to memorize content with and without emotional valence, either matching face pairs or Chinese character with unrelated German nouns for 3 weeks. This was followed by a period of relearning the already learnt, though shuffled then associative pairs, while subjects received daily escitalopram or placebo treatment. Although we hypothesized that SSRI intake, in combination with emotional reversal learning, would amplify sMRI-detectable grey matter changes in the hippocampus, parahippocampus, amygdala, prefrontal cortex, insular and the anterior cingulate cortex. We found no effects of escitalopram (vs. placebo) or relearning (as well as previous learning) on cortical and subcortical grey matter. To ensure methodological reliability, we pursued a surface-based and voxel-based grey matter analysis using surface-as well as voxel-based methods, where no pre-post substance or (re)learning interactions in brain structure were found.

Structural MRI has been used to examine learning-induced neuroplasticity in numerous intervention studies (Draganski, et al., 2004). As a consequence of training and learning, physical activity and perceptual stimulation grey matter expansions have been found in stimulus-specific brain regions, suggesting that high-frequent non-pharmacological interventions impact neuronal activation and further prime MRI-detectable morphological changes (Delon-Martin, et al., 2013; Draganski, et al., 2006; Maguire, et al., 2000). Serotonergic agents have been found to alter the grey and white matter as well as neurotransmitter levels (Hahn, et al., 2010; Kraus, et al., 2017; Seiger, et al., 2021; Sladky, et al., 2015; Spurny, et al., 2021), affect learning and cognition in healthy cohorts, while most importantly being designated as clinically relevant (Knorr, et al., 2019; Nissen, et al., 2010). With this study, we sought to facilitate neuronal plasticity and thus, enable adaptational processes in the brain that can be observed with MRI. Also, we applied associative learning with emotional content in comparison to semantic associative learning, which was intended to influence brain areas critical for emotional learning. In line, behavioral normalization following chronic fluoxetine administration was tightly linked to positive environmental influences and authors concluded that fluoxetine did not directly increase mood symptoms, but rather enables stimulus quality dependent neuronal reorganization (Alboni, et al., 2017). As a translational success, Branchi and colleagues further found similar effects of fluoxetine in patients with depression, where symptom trajectories during SSRI treatment were reliant on social environmental influence (Chiarotti, et al., 2017).

Learning and memory in general is ascribed to the hippocampus, the primary area for memory consolidation, which orchestrates inputs in close relation with other limbic regions and the prefrontal cortex (Pajkert, et al., 2017). Emotional processes in memory and extinction, where mostly fear is utilized as an emotional valence in experimental animal procedures, is mapped to brain regions like the amygdala, orbitofrontal cortex, hippocampus (Herry and Johansen, 2014). Dysfunctional limbic networks and abnormal monoaminergic signaling are associated to negative affective states, whereas successful treatment with SSRIs has been found to normalize attentional bias (Castren, 2005; Murray, et al., 2011; Zhou, et al., 2015). When testing for the interaction effects drug intervention and relearning on brain structure, we observed no differences in any emotion-processing brain regions included in our hypothesis. In a second step, we included all cortical regions in one model and volumes of subcortical brain regions in a 2^nd^ model and again found no influence of substance. In a multimodal imaging study that aimed to investigate age-dependent plastic effects in the visual cortex following texture training, white matter changes were found only in elderly, with apparent functional, but no structural differences in the visual cortex after training in younger individuals. Authors suggest that the brains of elderly needed to undergo structural changes were to increase performance, whereas altered neuronal activation in younger individuals was sufficient to raise performance levels (Yotsumoto, et al., 2014). As our sample of healthy individuals was comparably young with an average age of 25, it is possible that the learning intervention did cause changes on a neuronal level, though rather on an activity or functional level than on brain structure (Yotsumoto, et al., 2014). Functional and metabolic studies, investigating task-based neuronal activity as well as glutamate and GABA levels after relearning and escitalopram administration, found escitalopram-and relearning-driven interactions in different brain regions (Reed, et al., 2021; Spurny, et al., 2021). Thus, time interval and intensity of interventions as applied in this study might be extensive enough to alter activity and metabolic measures, though are not sufficient to affect brain structure. Also, paradigm parameters of this study might have been fallen short to induce structural changes. Quantity of learning (e.g., daily time spent with the task, extent of scheduled days) might just not have sufficiently provoked alterations of morphological measures as cortical thickness and subcortical volumes. Furthermore, even small doses of the SSRI citalopram led to substantial serotonin transporter occupancy and thus, assumed to initiate the drug’s pharmacodynamical potency (Meyer, et al., 2004). In our study, citalopram blood levels were beneath the reference area for escitalopram in half of the individuals that received escitalopram, according to (Hiemke, et al., 2011). Participants received non escalated therapeutic doses of escitalopram, to prevent higher drop-out rates due to tolerability issues that correspond to higher dosing (Furukawa, et al., 2019). Nevertheless, we cannot exclude that higher doses of escitalopram or other SSRIs would have instigated detectable grey matter differences. An abundance of literature, from pharmacological to genetic and (pre)clinical studies, aimed to disclose the involvement of serotonin in neurobiological mechanisms and their relevancy to neuropsychiatric disorders. Given this vast effort in the field of neuroimaging, studies on the impact of serotonergic agents on grey matter *in vivo* using MRI are surprisingly rare. In a placebo-controlled study in monkeys, sertraline 20 mg/kg was applied for 18 months (this equals approximately 5 human years), a frequently used SSRI, was found to decrease the volume of the hippocampus and anterior cingulate cortex. Contrary, in depressed and SSRI-treated monkeys analyses revealed distinct trajectories of grey matter changes, describing enhancements in the bilateral hippocampus and the left anterior cingulate (Shively, et al., 2017). In a voxel-wise structural study by our group in healthy humans with smaller sample size, 10 days of escitalopram was related to increased grey matter of multiple brain regions including the posterior cingulate cortex, ventral precuneus, the insula and the fusiform gyrus, while functional connectivity correlated with structural findings (Kraus, et al., 2014). Functional MRI studies repeatedly describe neuronal activation in various brain regions subsequent to learning and drug intervention. Thus, we expected that SSRI-or learning-induced functional activation would provoke structural changes. Contrary to previous findings, we found neither grey matter changes after a period of associative learning nor a succeeding period of relearning in combination with and without daily SSRI intake. Findings suggests that SSRI intake or learning, as applied in this study setting, have only limited impact on structural changes measurable with MRI using a voxel size of 1.12mm^3^ and surface-and voxel-based measures.

## Limitations

Even though, this study boasted a rather large sample and state of the art methods were used to analyze the data this longitudinal study has limitations that may compromise the interpretation of its results. Since we did not observe differences in interactions between learning groups and time regarding its effects on brain structure, it is possible that the chosen time span of 7 minutes learning daily for several weeks was chosen too sparsely. Further, a limitation could be that the T1-weigthed imaging with a voxel size of 1.1 mm^3^ processed via automatic segmentation pipelines for the measurement of cortical thickness and subcortical volumes was too unsusceptible to detect these changes, but also VBM analysis did yield no results. Even though, the SSRIs plasma levels of all subjects were measured at 3 time points throughout the relearning phase, we cannot be certain that all subjects were fully compliant and regularly took their daily oral dose over 3 weeks.

## Conclusion

We found that subacute SSRI administration by associative relearning interaction significantly modulate cortical thickness and subcortical volumes in healthy subjects. We cannot exclude that the dosing of escitalopram (although therapeutical doses were administered) or the intensity and duration of learning and relearning was designed to sufficiently to induce grey matter changes in healthy subjects. In previous studies functional and neurotransmitter changes have been found following to SSRI and relearning, whereas here, time interval and intensity of interventions might not have been ample enough to affect brain structure. We expected greater and wider effects of escitalopram and relearning on the brain structure. However, further investigations will have to examine SSRIs effects on relearning and structural neuroplasticity.

## Acknowledgements

We wish to thank the team members as well as the medical students of the Neuroimaging Lab (NIL), headed by Prof. Lanzenberger, the corresponding author.

## Declarations

### (i) Funding

The analysis is part of a larger study that was supported by the Austrian Science Fund (FWF) grant number KLI 516 to R.L., the Medical Imaging Cluster of the Medical University of Vienna, and by the grant "Interdisciplinary translational brain research cluster (ITHC) with highfield MR” from the Federal Ministry of Science, Research and Economy (BMWFW), Austria. M. K. is recipient of a DOC-fellowship of the Austrian Academy of Sciences at the Department of Psychiatry and Psychotherapy, Medical University of Vienna. D.P. is supported by the MDPhD Excellence Program of the Medical University of Vienna.

### (ii) Conflicts of Interest

There are no conflicts of interest to declare regarding the present study. R. Lanzenberger received travel grants and/or conference speaker honoraria within the last three years from Bruker BioSpin MR and support from Siemens Healthcare regarding clinical research using PET/MR. He is a shareholder of the start-up company BM Health GmbH since 2019. S. Kasper has received grants/research support, consulting fees and/or honoraria within the last three years; grant/research support from Lundbeck; he has served as a consultant or on advisory boards Celegne, IQVIA, Janssen, Lundbeck, Mundipharma, Recordati, Takeda and Schwabe; and he has served on speakers bureaus for Angelini, Aspen Farmaceutica S.A., Janssen, Krka Pharma, Lundbeck, Medichem Pharmaceuticals Inc., Neuraxpharma, OM Pharma, Pierre Fabre, Sanofi, Servier, Schwabe, Sun Pharma. C. Kraus received travel grants from Roche and AOP Orphan Austria, speaker honoraria from Janssen.

### (iii) Availability of data and material

The full data can be made available upon reasonable request to the corresponding author.

